# *De Novo* Design of Anti-Aging Peptides

**DOI:** 10.1101/2025.10.04.680435

**Authors:** Yan Pan, Li Fan, Nan Zhang, Jiexin Zheng, Zhong Jin, Hongxia Cai, Haomiao Ma, Wenjing Zhang, Jingyuan Zhu, Wentao Xu, Yiwen Gong, Ruofei Li, Rui Liang, Guojun Li, Haiming Jing, Junyu Ning, Shan Gao, Bo Xian

## Abstract

Aging, a universal biological process in complex organisms, is increasingly recognized to be driven by progressive loss of epigenetic information, as proposed in the Information Theory of Aging (ITOA). However, research on anti-aging peptides remains scarce, with most existing efforts confined to derivatives of natural proteins, while systematic design attempts are virtually absent. This limitation not only restricts discovery within the evolutionary sequence space but also hampers the identification of candidates with novel mechanisms and improved efficacy. Here, we present ElixirSeeker2, the first computational framework for *de novo* design of anti-aging peptides. By integrating modeling of known anti-aging peptides, activity scoring from the IC50 database, and penalty constraints from toxic peptides, ElixirSeeker2 enables large-scale virtual screening and identification of novel peptide candidates. Several lead peptides demonstrated significant effects in delaying cellular senescence, restoring cellular functions in vitro and in enhancing locomotor activity of aged *Caenorhabditis elegans*. This study not only validates the feasibility of de novo design in anti-aging interventions but also establishes a strategy for the development of next-generation biologics.

## 1. Introduction

Aging is an inevitable biological process in all complex organisms, characterized by the progressive decline of physiological functions and increased susceptibility to multiple diseases, including neurodegenerative disorders, cardiovascular diseases, and cancer [1–3]. Despite its complexity and heterogeneous manifestations, researchers have long sought to identify a unifying mechanism that drives aging. Recently, an information-theoretic framework has emerged as a novel perspective. All analog information storage systems, including biological systems, share a fundamental weakness: they are inherently susceptible to noise, which gradually erodes or distorts the original information [4].

In biology, this concept is exemplified by the dynamic alterations of the epigenetic landscape. The epigenome, as a central repository of information guiding gene expression programs, is essential for maintaining cellular identity and function. However, over time, persistent challenges such as cell division, environmental stress, and endogenous damage progressively disrupt and erode epigenetic information. This process has been demonstrated as a major driver of cellular aging in both yeast and mammalian models [5–7].

Building on this evidence, Sinclair and colleagues proposed the Information Theory of Aging (ITOA) [8], which posits that aging is fundamentally a process of information loss, particularly the gradual erosion of epigenetic information. Crucially, studies have shown that restoring information integrity via epigenetic reprogramming can significantly reverse cellular aging phenotypes and restore tissue functions, highlighting remarkable regenerative potential [9–11]. Thus, ITOA provides not only a unifying theoretical framework for aging but also a guiding principle for novel anti-aging interventions.

According to ITOA, effective strategies to delay or reverse aging should focus on “supplementing” or “stabilizing” critical cellular information. In this context, proteins—the primary executors of life’s functions—naturally emerge as key therapeutic targets. Numerous studies have identified proteins that play central roles in aging regulation. For example, Klotho, one of the earliest identified aging-suppressor proteins, significantly extends lifespan when overexpressed in mice, whereas its deficiency leads to premature aging syndromes [12]. Similarly, members of the SIRT family and FOXO transcription factors are critically involved in lifespan regulation, metabolic homeostasis, and stress resistance [13–15].

These discoveries raise a pivotal question: can proteins themselves serve as “information carriers” that, when supplemented exogenously, delay or reverse the loss of cellular information? Compared with large full-length proteins, short peptides (typically fewer than 50 amino acids) offer several advantages as therapeutic agents: smaller molecular weight, ease of synthesis and modification, high tissue penetrability, and low immunogenicity [16–18]. In recent years, functional peptide fragments derived from natural anti-aging proteins have been identified. For instance, Klotho-derived peptides have demonstrated protective effects on cognitive function and kidney health [19].

Despite these advances, most current anti-aging peptide development remains restricted to derivatives of natural sequences. While such strategies can yield biologically active candidates, they are confined within the sequence space shaped by natural evolution, limiting the discovery of molecules with novel mechanisms or superior efficacy [20]. In small-molecule drug discovery, high-throughput phenotypic screening and machine learning approaches have already facilitated the identification of novel anti-aging compounds such as ElixirSeeker [21–22]. Likewise, advances in artificial intelligence, particularly deep learning, have enabled powerful modeling of protein structure and function, making *de novo* peptide and protein design increasingly feasible [23–25].

Compared to traditional derivative strategies, *de novo* design offers substantial advantages. First, it does not rely on any known protein templates, thereby enabling exploration of a vastly broader sequence and structural space, which may yield entirely novel peptides not found in nature. Second, *de novo* design allows precise tuning of physicochemical properties, which is essential for optimizing drug-like features such as enhanced cell penetration, improved serum stability, and reduced toxicity [26].

In this study, guided by the Information Theory of Aging, we present ElixirSeeker2, a novel framework for *de novo* design of anti-aging peptides. Our approach integrates modeling of existing anti-aging peptides, activity scoring using the IC50 database, and penalty terms from toxic peptide datasets to enforce safety constraints as shown in Figure 1. Through large-scale virtual screening, we identified lead peptide candidates capable of delaying cellular senescence and restoring cellular functions. Importantly, our work not only provides a collection of entirely new anti-aging peptide candidates but also validates the feasibility and great potential of *de novo* design as a paradigm-shifting strategy for the development of anti-aging biologics.

**Figure 1.**
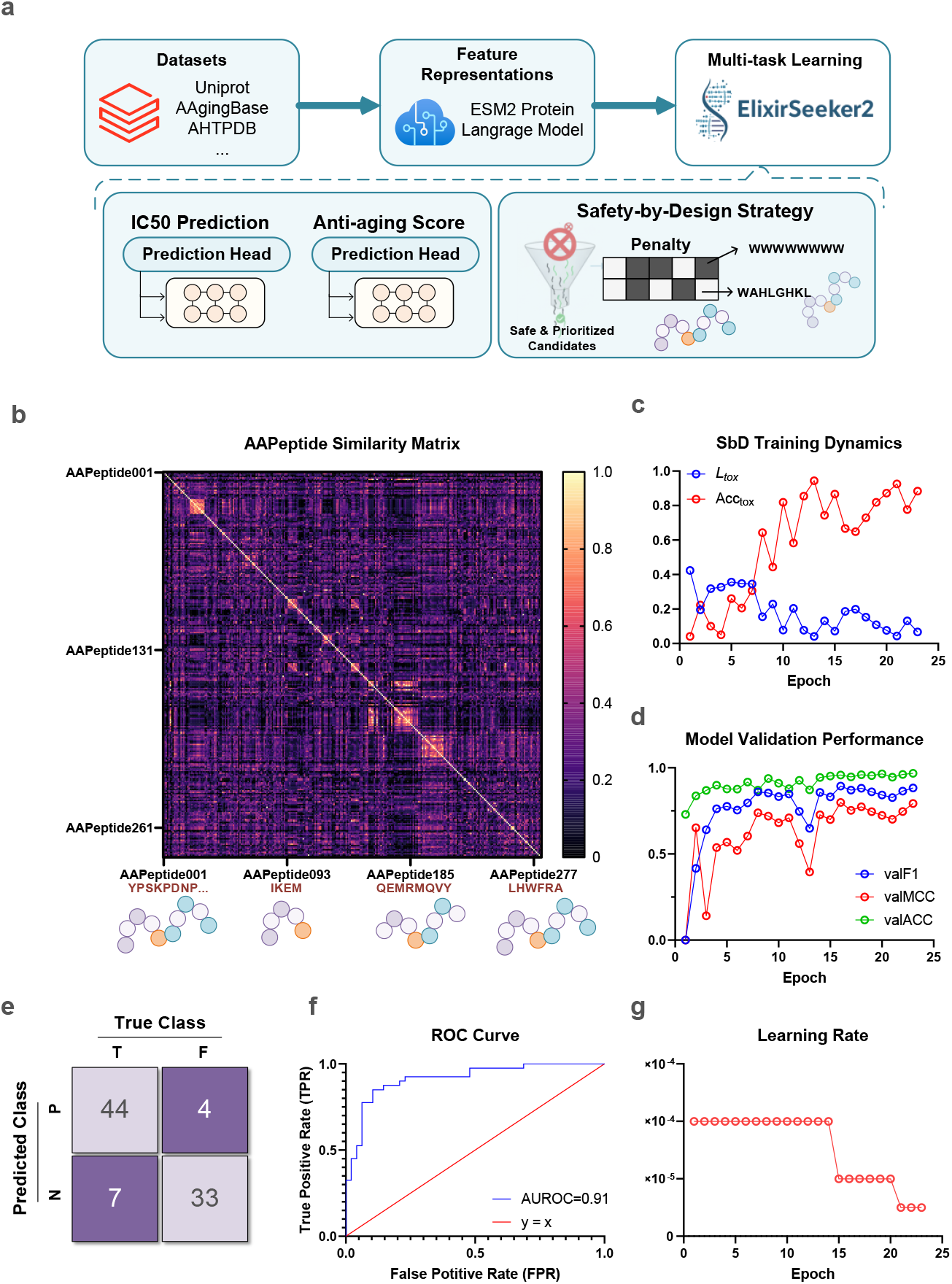
The proposed dual-head multi-task learning model of *de novo* anti-aging peptide design and its validation performance. (**a)** The classification branch provides an anti-aging score by distinguishing positive samples from AgingBase (experimentally validated anti-aging peptides) and negative counterparts extracted from UniProt with matched sequence lengths. In parallel, the regression branch leverages IC_50_-annotated peptides from AHTPDB to model general bioactivity. A Safety-by-Design strategy is further integrated by incorporating known toxic peptides into the loss function as penalty terms, ensuring that predictions balance functional efficacy with safety considerations. **(b)** Similarity matrix of anti-aging peptides (AAPeptides) encoded by ESM-2, where dark purple indicates low similarity and bright yellow indicates high similarity. **(c)** Loss_tox and Accuracy_tox during SbD strategy training. The decreasing toxicity penalty indicates stable and effective model training. **(d)** Performance metrics (F1, MCC, and ACC) of the anti-aging binary classification model on the validation set. **(e)** Confusion matrix of the anti-aging binary classification model on the validation set. **(f)** ROC curve of the anti-aging binary classification model with an area under the curve (AUC) of 0.91. **(g)** Dynamic change of learning rate across training epochs.

## 2. Results and Discussion

### 2.1. Model design rationale and validation performance

To address the scarcity and abstract nature of experimentally validated anti-aging peptides, we designed a dual-head multi-task learning framework under a Safety-by-Design (SbD) strategy (Figure 1a). Since the number of confirmed anti-aging peptides is limited, relying solely on classification is insufficient. Therefore, the architecture incorporates complementary learning objectives to capture multiple aspects of peptide functionality.

Specifically, the classification head provides an anti-aging score by discriminating validated peptides from AAgingBase against length-matched peptides sampled from UniProt. In parallel, the regression head leverages IC_50_ values from AHTPDB to encourage the model to recognize peptides with general biological potency. To further enforce safety, toxic peptides, including membrane-disruptive and hydrophobic motifs were integrated into the loss function through an additional penalty term, enabling the model to disentangle beneficial bioactivity from toxicity and thus balancing efficacy with safety.

As shown in Figure 1b, the similarity matrix of anti-aging peptides (AAPeptides) encoded by ESM-2 demonstrates that the majority of peptide pairs exhibit low similarity, indicating both the intrinsic diversity of the dataset and the difficulty of identifying universal sequence patterns. This diversity also ensures that the model avoids overfitting to narrow sequence motifs. Importantly, we deliberately adopted ESM-2 embeddings without designing overly complex architectures, given three considerations: (i) the pretrained model already provides sufficiently rich representations of peptide sequences; (ii) the majority of anti-aging peptides are short (6–25 amino acids), reducing the necessity of deep and computationally expensive models; and (iii) a more parsimonious architecture facilitates interpretability and generalizability.

To evaluate the effectiveness of the SbD strategy, we monitored the toxicity-related loss (Loss_tox) and toxicity classification accuracy (ACC_tox) during training (Figure 1c). toxicity-related loss consistently decreased across epochs, while toxicity classification accuracy increased, suggesting that the model progressively learned to identify toxic motifs. From a drug discovery perspective, the rising toxicity classification accuracy further validates the ability of the framework to distinguish toxic from non-toxic peptides, confirming that the SbD mechanism successfully decouples functional motifs from toxicity-related patterns within the latent space. Such safety-aware learning is particularly critical for de novo peptide design, where preclinical toxicity remains a major bottleneck in translational applications.

Model validation results further confirmed performance. After 25 epochs, early stopping was triggered (Figure 1g). The classification model achieved high performance with F1 and MCC values exceeding 0.85 and an accuracy (ACC) approaching 0.90 (Figure 1d). The confusion matrix (Figure 1e) highlights balanced predictive power across positive and negative classes, while the receiver operating characteristic (ROC) curve yielded an AUROC of 0.91 (Figure 1f), underscoring discriminative capability. The dynamic adjustment of the learning rate (Figure 1g) demonstrates stable convergence within relatively few epochs, reflecting the sufficiency of the training data and the efficiency of the model architecture.

In summary, the proposed framework effectively integrates functional, safety, and general bioactivity constraints, yielding a robust predictive model for anti-aging peptides while ensuring safety, a critical requirement for subsequent *de novo* peptide design.

### 2.2. Large-scale *de novo* peptide screening and candidate characterization

To systematically explore the anti-aging peptide sequence space, we adopted two complementary strategies to generate candidate sequences (Figure 2a). For 6-mer peptides, we performed an enumeration of the entire combinatorial space (20^6^ ≈ 64 million). For longer peptides ranging from 7 to 25 amino acids, we employed a genetic algorithm to efficiently sample approximately 10 million peptides for each length, resulting in ∼251 million sequences in total.

**Figure 2.**
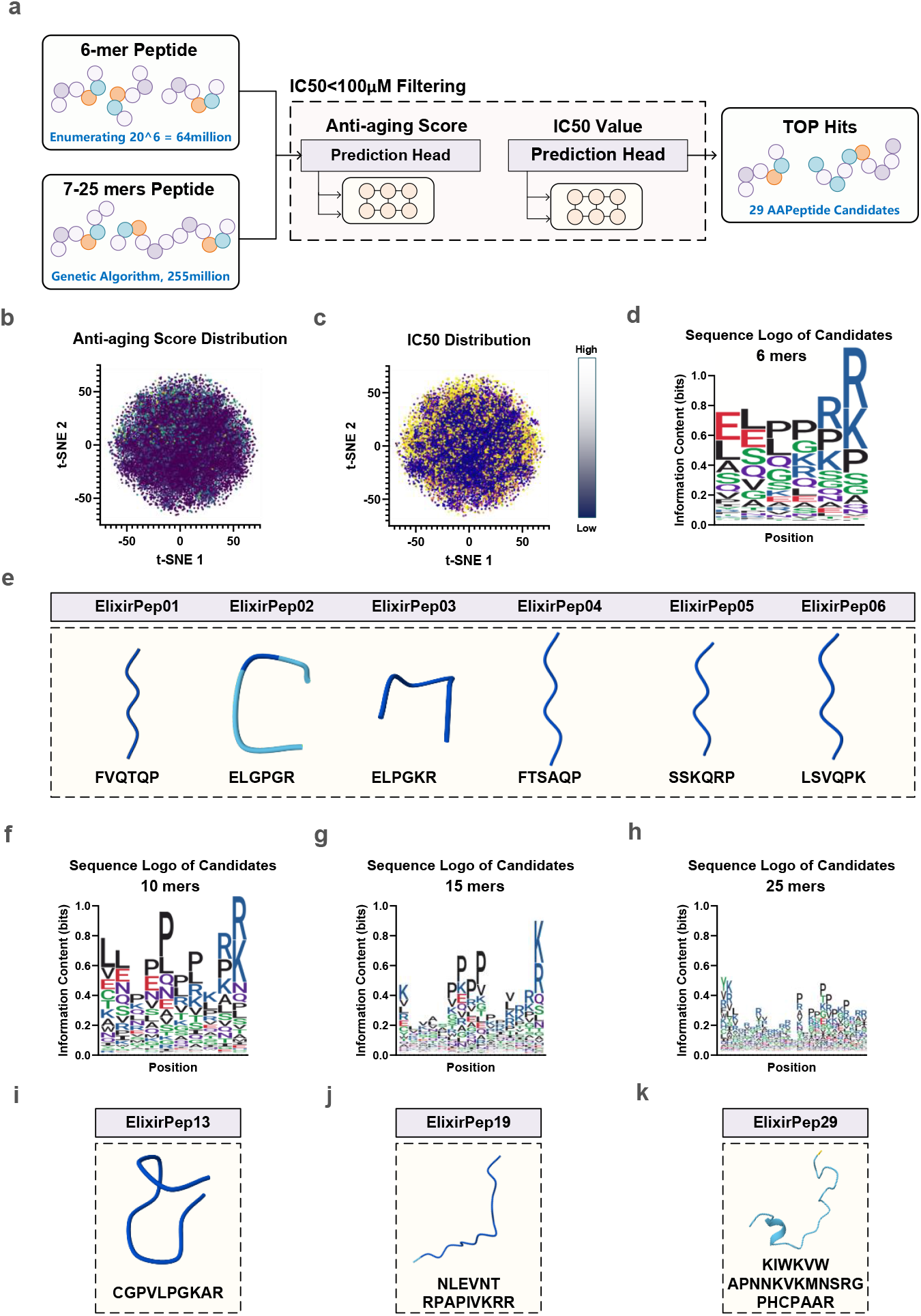
Large-scale peptide screening and sequence characterization. **(a)** Schematic overview of candidate generation: exhaustive enumeration of all 6-mer peptides (∼64 million) and genetic algorithm sampling of 7–25-mer peptides (∼251 million). After IC_50_ < 100 μM filtering, ∼200 million peptides were evaluated by the predictive framework to yield 29 top-ranked candidates. **(b)** t-SNE projection of anti-aging score distribution for filtered peptides **(c)** t-SNE projection of IC_50_ distributions. **(d)** Sequence logo of 6-mer candidate peptides **(e)** Examples of top 6-mer peptides (ElixirPep01–06) with sequences and predicted 3D conformations **(f–h)** Sequence logos for candidate peptides of 10, 15, and 25 residues, respectively, showing recurrent enrichment of R/K/P residues at the C-terminus. **(i–k)** Representative top-ranked peptides of 10, 15, and 25 residues (ElixirPep13, ElixirPep19, and ElixirPep29). For the all seqlogos, see Figure S1.

This approach was chosen to decouple the predictive model from the generative process, thereby avoiding reliance on generative AI models, which often impose strong distributional priors and risk collapsing into biased sequence motifs. Instead, the enumeration and genetic search ensured a maximally diverse candidate pool that allowed the predictive model to operate in a hypothesis-driven, unbiased manner.

The initial sequence pool of ∼315 million peptides was filtered by predicted IC_50_ values, retaining only those with IC_50_ < 100 μM as a threshold for potential biological activity. This yielded ∼200 million peptides for further screening. Figure 2b illustrates the distribution of anti-aging scores across this filtered set in t-SNE space, where the majority of sequences exhibited very low predicted scores, highlighting the rarity of potential anti-aging candidates. Interestingly, when comparing the IC_50_ distribution (Figure 2c), we observed that peptides predicted with higher IC_50_ values often coincided with elevated anti-aging scores, whereas peptides with low IC_50_ values (indicative of strong bioactivity) generally had low anti-aging scores. This suggests that while general potency is necessary, anti-aging functionality may be associated with milder or more selective activity rather than indiscriminate high potency, a property that could translate into better tolerability and lower off-target effects.

Sequence logo analysis of top-scoring peptides provided further mechanistic insight. For 6-mer peptides (Figure 2d), the most enriched motif was ELPPRR; however, when examining the individually highest-ranked sequences (ElixirPep01–06, Figure 2e), only limited overlap with the logo was observed (e.g., FVQTQP, ELPGKR). This discrepancy reflects the fact that sequence logos represent averaged positional preferences across the candidate pool, whereas the model predictions capture context-dependent interactions between residues. In other words, the model does not simply reward isolated motifs but instead evaluates their contribution within the global structural and biochemical context.

Similarly, sequence logos for longer peptides (10, 15, and 25 residues) revealed consistent enrichment of C-terminal arginine/lysine/proline residues (Figures 2f–h), yet the top individual peptides (ElixirPep13, ElixirPep19, and ElixirPep29; Figures 2i–k) again displayed distinctive local features, suggesting the context-dependent nature of functional predictions.

Notably, the recurrent enrichment of R/K/P residues at the C-terminus has strong biological plausibility: (i) the positive charges of arginine/lysine facilitate interaction with negatively charged phospholipids in cell membranes, a hallmark of cell-penetrating peptides; and (ii) R/K/P residues at terminal positions are unfavorable cleavage sites for canonical endopeptidases, thereby conferring enhanced proteolytic stability. Together, these features suggest that the model has implicitly captured hallmarks of membrane penetration and in vivo stability.

Another recurrent pattern was the frequent appearance of proline or consecutive prolines (PP), which are well known as α-helix breakers and structural inflection points that induce β-turns or folding interruptions. Such motifs are consistent with the presence of modular architectures in short peptides, where one segment anchors or penetrates the membrane, and the other executes the functional activity. This observation suggests that the model not only identifies membrane-active features but also recognizes design principles for bifunctional peptide modules.

Overall, these analyses highlight the capacity of the framework to generate and prioritize candidates with both mechanistic plausibility and novel sequence diversity, while maintaining safety and functional considerations.

### 2.3. Proline as a position-dependent hallmark of anti-aging peptides

In Section 2.2, we performed self-normalized enrichment analysis, where candidate peptides were compared internally to highlight recurrent sequence patterns. While informative, this approach does not account for biases in the natural sequence space. To address this, we further implemented a length-matched position-specific residue enrichment analysis, using peptides extracted from UniProt as a background. For each peptide length between 6 and 25 residues, the top 1,000 sequences ranked by anti-aging score were selected as candidates, and their amino acid frequencies at each position were compared against the background distribution.

Fold-change (FC) values were calculated along with false discovery rate (FDR) corrections, allowing the identification of significantly enriched (red) or depleted (blue) residues with thresholds of FDR < 0.05 and |FC| > 2.

As shown in Figure 3a, volcano plots revealed systematic amino acid biases across peptide lengths. Proline (P) emerged as the most significantly enriched residue overall, consistent with our findings in Section 2.2. However, the enrichment of proline was not uniform across all positions. For example, Figure 3b illustrates position-wise analysis for 6-mer peptides, where proline was significantly depleted at position 2 (6,2,P), while enrichment was observed closer to the C-terminal region in longer peptides (e.g., 16,13,P). Similarly, several acidic residues such as aspartic acid (D) and glutamine (Q) were consistently depleted at specific positions (e.g., 12,8,D).

**Figure 3.**
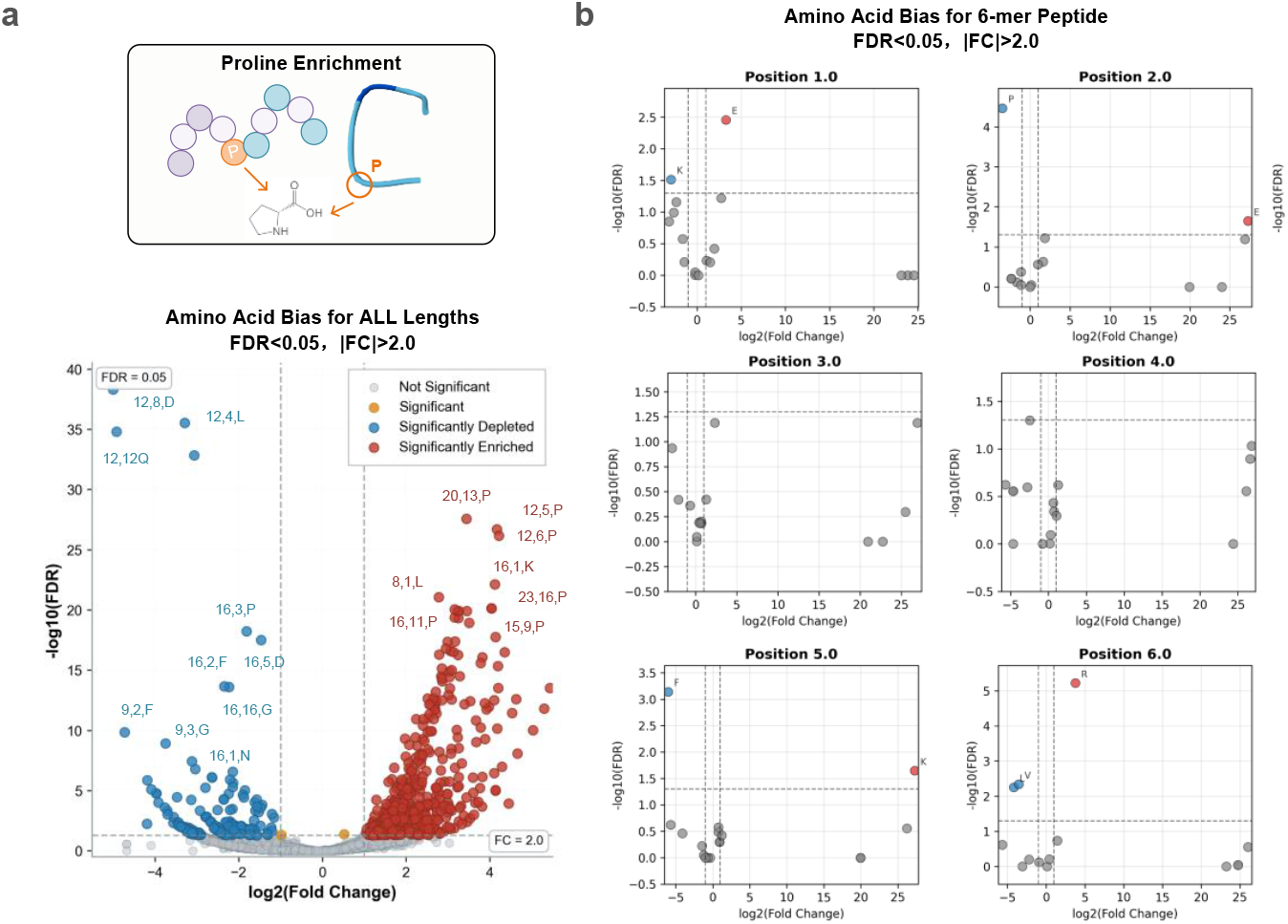
Position-specific residue enrichment analysis of anti-aging peptides. **(a)** Volcano plot of amino acid enrichment across candidate peptides compared with length-matched UniProt background sequences. Significantly enriched residues (red, FC > 2, FDR < 0.05) and depleted residues (blue, FC < −2, FDR < 0.05) are highlighted, with annotations indicating peptide length, position, and residue identity (e.g., “20,13,P” = position 13 of a 20-mer). **(b)** Position-specific analysis of 6-mer peptides, showing fold-change distributions at each position.

These results suggest that proline enrichment in anti-aging peptides is position-dependent. At the N-terminus, proline is often unfavorable, possibly because its conformational rigidity disrupts the initiation of stable secondary structures or hinders recognition motifs. In contrast, enrichment near the C-terminus likely reflects its structural role as a β-turn inducer or helix breaker, introducing conformational kinks that facilitate modular peptide architectures. Such C-terminal enrichment may further synergize with positively charged residues (R/K) frequently observed in this region (Section 2.2), collectively promoting both membrane interaction and proteolytic resistance.

Taken together, this length- and position-specific analysis confirms that the predictive model captures not only global compositional biases but also context-sensitive structural preferences, highlighting proline as a key determinant of anti-aging peptide architecture when positioned appropriately within the sequence.

### 2.4. Candidate peptides effectively delay or reverse cellular and organismal aging

Given that all 6-mer peptides were exhaustively enumerated in silico, their likelihood of success was expected to be relatively higher compared to longer peptides. Accordingly, six top-ranked 6-mers (ElixirPep01–06) were selected for synthesis and experimental validation, while 23 peptides from the 7–25-mer pool were chosen based on IC_50_ values, sequence diversity, and peptide length. In total, 24 peptides were successfully synthesized at >95% purity (Table S1).

To evaluate their anti-aging activity, we implemented two treatment strategies in HEK293 cells (Figures 4a and 4e). Although HEK293 is an immortalized cell line, recent studies have demonstrated its suitability as a senescence model due to robust stress-induced phenotypes and high experimental efficiency[27-29]. The first approach was co-treatment, in which peptides were administered concurrently with H_2_O_2_-induced senescence, while the second was post-treatment, in which peptides were applied after senescence induction to assess their capacity for phenotype reversal.

**Figure 4.**
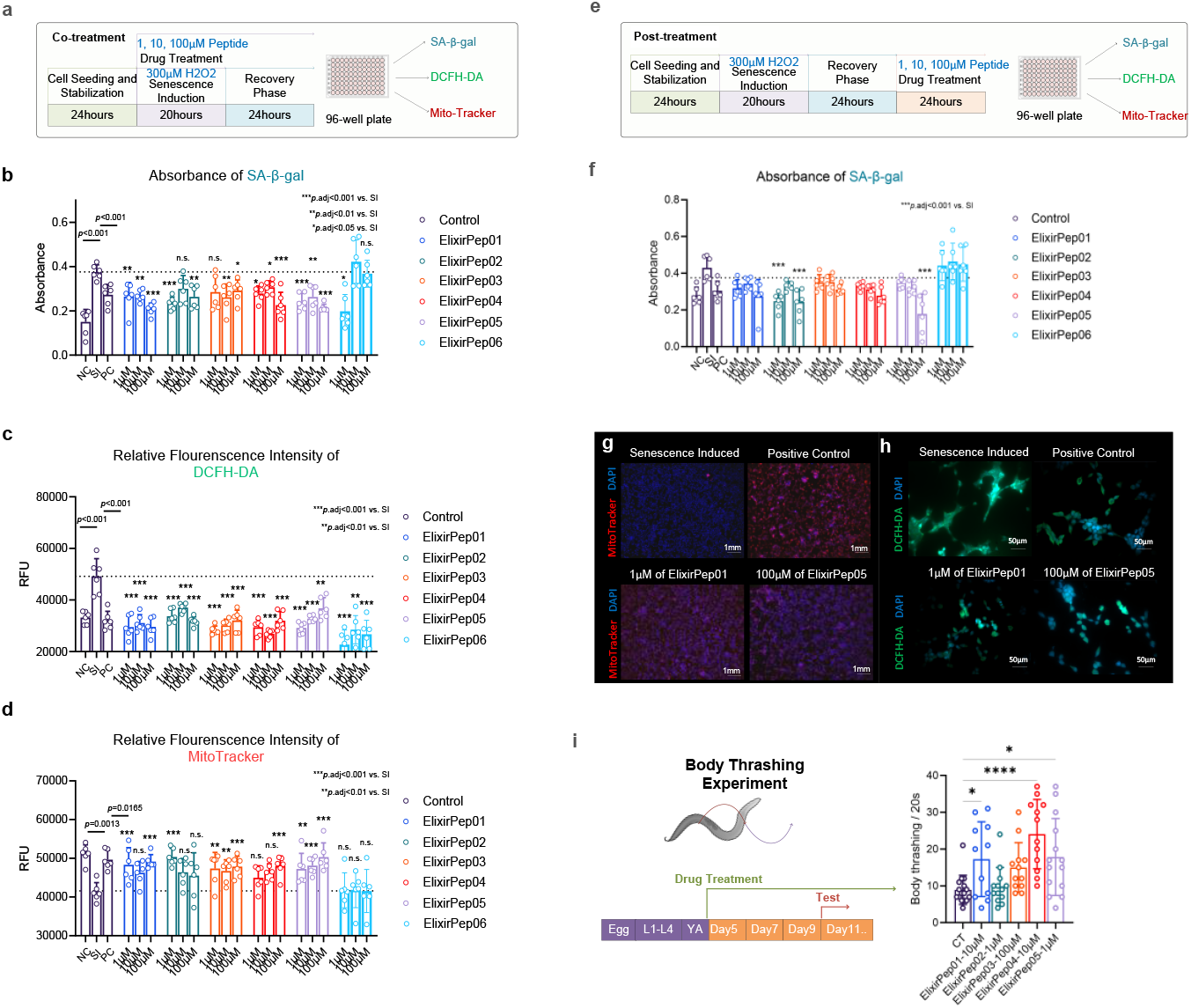
Experimental validation of candidate peptides in cellular and organismal models of aging. **(a)** Co-treatment paradigm: peptides were applied concurrently with H_2_O_2_-induced senescence, followed by assessment of SA-β-gal, DCFH-DA, and MitoTracker signals. **(b–d)** Co-treatment results in HEK293 cells: SA-β-gal absorbance **(b)**, intracellular ROS levels **(c)**, and mitochondrial membrane potential **(d). (e)** Post-treatment paradigm: peptides were applied after senescence induction. **(f)** SA-β-gal absorbance in post-treatment assays. **(g–h)** Representative microscopy of ElixirPep01 and ElixirPep05 treatments: DAPI/MitoTracker (g, red = mitochondrial potential) and DAPI/DCFH-DA (h, green = ROS). Both peptides improved mitochondrial integrity and reduced ROS. **(i)** *C. elegans* body thrashing assays. For panels (b–d, f), data represent mean ± SEM of biological replicates. Statistical significance was assessed by two-way ANOVA with Benjamini–Krieger–Yekutieli FDR correction (q < 0.05) for post hoc comparisons versus SI group. For panel (i), body thrashing frequency was compared across groups by two-way ANOVA with the same post hoc correction versus CT group. *p < 0.05, **p < 0.01, ***p < 0.001.

Across both designs, cells were exposed to three concentrations (1, 10, 100 μM) of each peptide, alongside non-treated controls (NC), senescence-induced controls (SI), and a positive control (PC, 100 μM NAC). A total of 75 groups were screened in 384-well plates. Following treatment, senescence markers were quantified, including SA-β-gal activity, intracellular ROS (DCFH-DA), and mitochondrial membrane potential (MitoTracker).

In the co-treatment paradigm, all six 6-mers demonstrated measurable effects. SA-β-gal staining revealed significant reductions for ElixirPep01–05 at multiple doses, while ElixirPep06 showed efficacy only at 1 μM (Figure 4b). ROS levels were consistently suppressed across all six peptides (Figure 4c), and mitochondrial membrane potential was significantly improved by ElixirPep01–05 (Figure 4d). These findings confirmed protective activity during senescence induction.

To rigorously test the ability to reverse aging phenotypes, post-treatment assays were conducted (Figure 4e). As shown in Figure 4f, most peptides reduced SA-β-gal activity relative to SI controls, with ElixirPep02 and ElixirPep05 achieving particularly strong effects at 100 μM (p < 0.001). Representative microscopy further confirmed these results: ElixirPep01 and ElixirPep05 significantly reduced ROS (green, DCFH-DA) and improved mitochondrial membrane potential (red, MitoTracker) compared to SI controls (Figures 4g–h).

Finally, functional validation was extended to *Caenorhabditis elegans* (*C. elegans*). Worms were continuously administered peptides from the young adult (YA) stage until day 11. Body thrashing assays—a classic readout of muscle vitality in aging worms—revealed that ElixirPep01 (10 μM, p < 0.05), ElixirPep04 (10 μM, p < 0.0001), and ElixirPep05 (1 μM, p < 0.05) significantly enhanced locomotor activity compared to controls (Figure 4i).

Collectively, these results demonstrate that at least three of the top six 6-mer peptides effectively delay or reverse cellular and organismal aging phenotypes. Among the 24 peptides tested, ∼40% exhibited measurable efficacy in reversing senescence in post-treatment assays (Table 1), with several surpassing the performance of the positive control NAC.

**Table 1.**
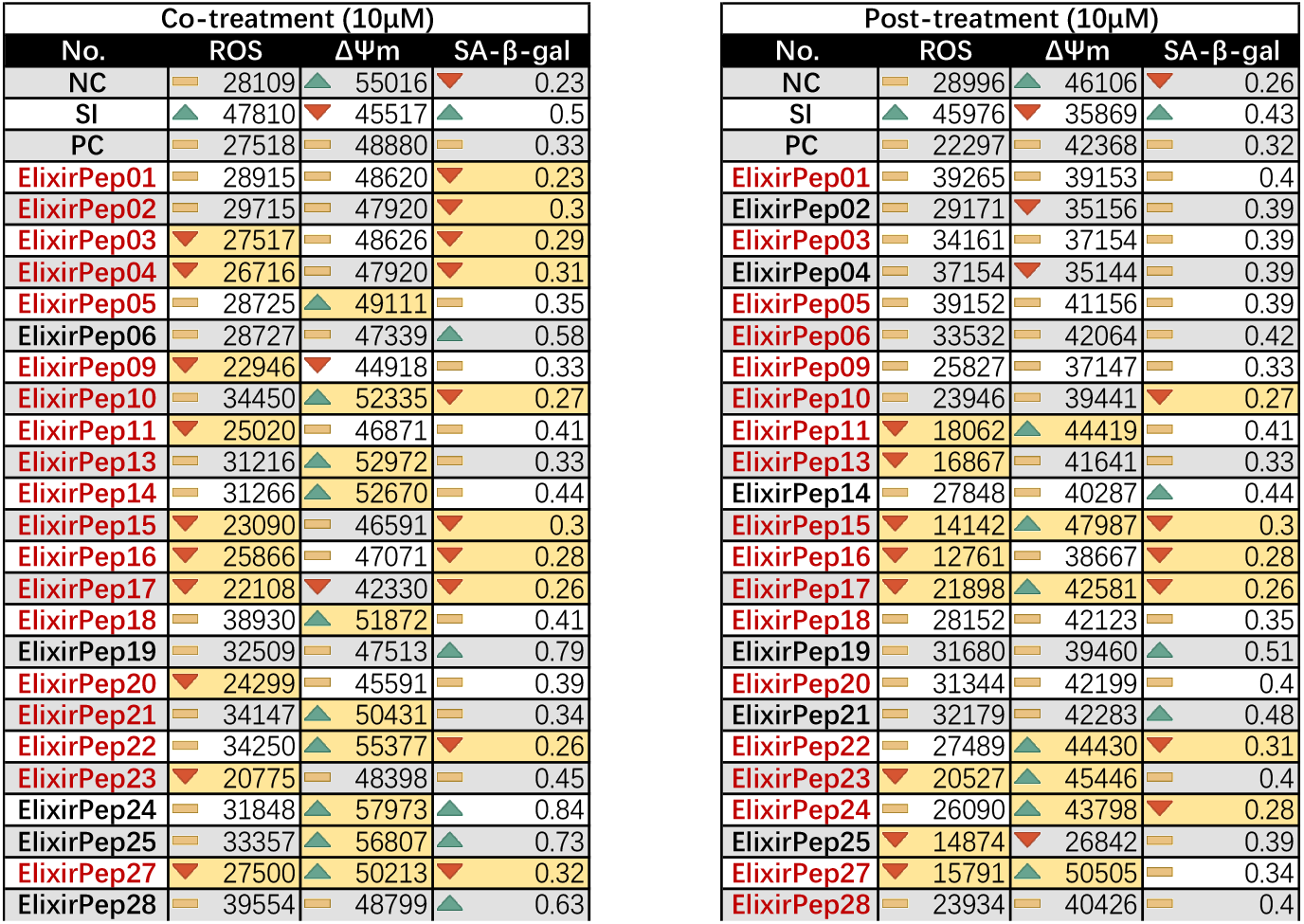
Effects of candidate peptides (10 μM) on cellular senescence markers in HEK293 cells under co-treatment and post-treatment conditions. Peptides with lower SA-β-gal absorbance than SI group and effective for both ROS and ΔΨm will be highlighted in red. Peptides with any one of these indicators exceeding the positive control level are highlighted in yellow.

## 3. Methods

### 3.1. Data and Multi-task Learning Framework Design

The present study employs a multi-task learning framework that integrates heterogeneous peptide datasets to construct a unified predictive model for anti-aging peptide discovery.

The central rationale is that experimentally validated anti-aging peptides are extremely scarce, limiting the effectiveness of conventional single-task classifiers. To mitigate this limitation, we deliberately introduce an auxiliary regression task that leverages peptide bioactivity data (IC50 values) not directly related to aging but reflective of general biological potency.

By jointly training on both classification and regression tasks within a shared representation space, the model is encouraged to capture broader biochemical and structural determinants of activity, thereby enhancing its ability to generalize to novel sequences with anti-aging potential.

For the classification task, positive samples were collected from AgingBase, a curated database of experimentally verified anti-aging peptides. These sequences represent the benchmark functional group of interest. As negative counterparts, we extracted peptide fragments ranging from 6 to 25 amino acids in length from the UniProt database. By contrasting AgingBase peptides with a diverse set of non-annotated short peptides from UniProt, the classification task is designed to highlight functional motifs and sequence-level properties associated with anti-aging activity.

To complement the classification task, a regression objective was constructed using peptides annotated with half-maximal inhibitory concentration (IC50) values obtained from the AHTPDB web server. Importantly, this dataset is not directly associated with anti-aging mechanisms, but rather with general peptide bioactivity in different biological contexts. The underlying assumption is not that IC50 serves as a surrogate marker of anti-aging efficacy, but that peptides with measurable activity embody structural and compositional features characteristic of functional biomolecules. By training the model to approximate IC50 values, the shared encoder is guided to learn gradients of bioactivity beyond binary decision boundaries.

Both tasks share a common sequence encoder derived from the Evolutionary Scale Modeling (ESM) framework, a large-scale protein language model pre-trained on millions of natural protein sequences. ESM embeddings capture contextualized residue-level dependencies informed by evolutionary constraints and structural regularities. By using ESM as the backbone, our approach benefits from transfer learning, where general protein-level representations are fine-tuned to the specific domain of short bioactive peptides. On top of the shared encoder, we implemented task-specific prediction heads: a fully connected layer with a sigmoid activation for binary classification of anti-aging versus non-anti-aging peptides, and a linear regression head for predicting IC50 values. Given the relative scarcity of anti-aging peptides compared to readily available negative samples, we adjusted the positive-to-negative ratio to approximately 1:1.2 to prevent strong class imbalance. Positive sequences were further augmented through controlled mutational perturbations, thereby enriching the diversity of the anti-aging set and reducing the risk of overfitting to narrow sequence motifs. Additionally, extreme homogeneous sequences were incorporated as contrastive examples, forcing the model to distinguish genuine biological signals from trivial repetitive patterns.

In summary, the data and model design of this study follow a principled strategy aimed at overcoming the challenges of limited positive samples and the risk of spurious correlations.

### 3.2. Loss Function and Safety-by-Design (SbD) Strategy

In addition to the multi-task learning framework, we further incorporated a Safety-by-Design (SbD) strategy to explicitly account for peptide toxicity. Toxicity represents one of the most critical barriers in the clinical translation of peptide-based therapeutics, and therefore cannot be neglected when building predictive frameworks for drug discovery. Our approach integrates prior knowledge of known toxic short peptides directly into the training objective by introducing a penalty mechanism: whenever the model assigns high anti-aging probability or strong activity scores to sequences documented as toxic, an additional loss term is applied to penalize such misclassifications. The rationale behind adopting SbD principles is threefold. First, biological functionality and toxicity are often entangled at the sequence level, and conventional activity-based tasks cannot guarantee safety. By incorporating explicit penalty signals, the model is encouraged to disentangle functional motifs from toxicity-associated patterns in its latent space. Second, SbD enhances the translational relevance of the framework by discouraging false optima—sequences that are predicted to be highly active yet unusable due to toxicity. Third, this strategy provides a safeguard for downstream generative tasks, ensuring that de novo designed peptides are guided not only by efficacy but also by inherent safety boundaries.

The overall loss integrates three components: an anti-aging classification loss, an activity regression loss, and a toxicity penalty term based on the principles of SbD. The total loss is defined as:

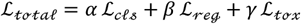

Where *α, β, γ* are hyperparameters balancing the contributions of different objectives.

The classification loss ℒ_*cls*_ adopts the Binary Cross-Entropy (BCE) function, ensuring discrimination between anti-aging and non-anti-aging peptides:

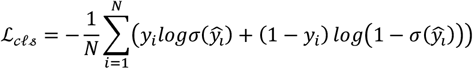

Here, *N* denotes the number of classification samples, *y*_*i*_ ∈ {0,1} is the ground-truth label, ŷ_*i*_ is the model logit, and *σ*(·) is the sigmoid activation.

The regression loss ℒ_𝓇ℯℊ_ is defined as the Mean Squared Error (MSE), guiding the model to capture continuous IC50 values:

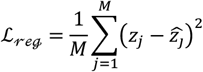

Where *M* is is the number of regression samples, *z*_*j*_ is the ground-truth IC50.

We further introduce the SbD-inspired toxicity penalty ℒ_*𝓉ℴ𝓍*_. Let ℒ denote the set of known toxic peptides. For each toxic sequence *k*, if the model incorrectly assigns a high anti-aging probability, an additional penalty is applied:

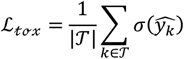

Where 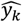 is the logit for toxic peptides. The penalty increases when 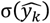 approaches 1 (misclassified as anti-aging) and vanishes when it approaches 0.

To further refine the control, we introduce a threshold parameter *τ*, penalizing only predictions above this safety margin:

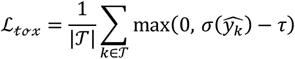

This formulation explicitly encodes avoidance of toxic misclassification into the training process, enforcing the model to separate functional from toxic sequence patterns in the latent space. To quantify the effectiveness of the SbD strategy, we used two key safety evaluation metrics: toxicity penalty and toxicity detection accuracy. The toxicity penalty is as described above as ℒ_*𝓉ℴ𝓍*_, and the toxicity detection accuracy is defined as

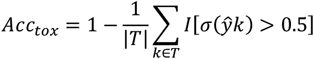

Where |*T*| represents the size of the toxic peptide set, and *I*[·] is an indicator function that returns 1 if the condition is true and 0 otherwise. *Acc*_*tox*_ it represents the proportion of toxic peptides correctly identified by the model as non-anti-aging peptides.

### 3.3. Model Optimization and scheduling

We used Adam (LR 1e-4), batch size 16, and up to 50 epochs. A ReduceLROnPlateau scheduler (factor=0.5, patience=3, min_lr=1e-6, cooldown=2) monitored validation total loss. Early stopping (patience=7, min_delta=0.001) restored the best weights upon convergence. Training ran on CUDA. Classification and regression minibatches alternate within each epoch, each backpropagating its respective loss term; the SbD penalty is attached to classification steps. After each epoch, we aggregate training losses and evaluate on the validation sets. The best checkpoint is saved with its configuration JSON.

We records all key metrics per epoch (total loss; task-wise losses; toxicity penalty; classification AUC/F1/MCC; regression RMSE/R^2^; LR history). We fixed random seeds (random_state=42) and standardized tokenization/length settings to improve reproducibility.

### 3.4. Compute Environment

All experiments were executed on a Linux workstation running Ubuntu 24.04 LTS, equipped with NVIDIA GeForce RTX 4090 accelerators. The cumulative compute budget for this study amounted to approximately 2000 RTX-4090 GPU-hours. In particular, the deep learning backbone was PyTorch 2.6.0 (paired with Triton 3.2.0 for fused kernels). The sequence modeling layer was built with Transformers 4.48.2, Tokenizers 0.21.0, and huggingface-hub 0.28.1. For scientific computing and numerical kernels we used NumPy 1.26.4 and SciPy 1.14.1, while classical metrics and model-selection utilities were provided by scikit-learn 1.6.1. Data wrangling and table-level persistence were implemented with pandas, and plotting/logging utilities relied on matplotlib 3.9.2, seaborn (matching our conda build), and tqdm (4.66+). On the CUDA side, our environment resolved to nvidia-cuda-runtime-cu12 12.4.127, nvidia-cudnn-cu12 9.1.0, nvidia-cublas-cu12 12.4.5.8, nvidia-nvjitlink-cu12 12.4.127, and related 12.4.x components. The Python toolchain was Python 3.12.

The full requirements file is shipped with the codebase.

### 3.5. Candidate Peptide Enumeration and Screening

To probe the peptide sequence space, we adopted a two-pronged strategy: exhaustive enumeration for 6-mers and genetic algorithm (GA)–based search for 7–25-mers. For hexapeptides, we enumerated the full combinatorial space over the 20 canonical amino acids.

All 64,000,000 6-mer sequences were scored by the trained multi-task model: we computed the anti-aging probability and the predicted IC50. Consistent with the activity prior, we filtered 6-mers using the threshold 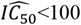 and ranked the survivors by anti-aging score.

For 7–25-mers, full enumeration is intractable. We therefore employed a standard GA to efficiently navigate the combinatorial landscape. For each length *L*∈{7.25}, we randomly initialized *N* mutually distinct sequences as the population, scored each sequence with the trained model, and iteratively evolved the population via selection, crossover, mutation, with elitism retained.

Across 6–25 aa, we evaluated 255,321,471 sequences in total. Within this subset, 19 distinct lengths (7–25 aa) were represented with the following counts: length 7—8,646,687, 8—8,990,820, 9—9,248,575, 10—9,351,083, 11—9,380,009, 12—9,508,744, 13—9,751,485, 14—9,961,420, 15—10,138,038, 16—10,502,810, 17—11,199,839, 18—11,814,377, 19—12,169,466, 20— 12,376,904, 21—12,490,351, 22—12,596,373, 23—12,696,600, 24—12,792,489, and 25— 12,858,105.

### 3.6. Peptide Synthesis and Preparation

Potential anti-senescence peptides identified from preliminary screening were synthesized by GenScript with a purity of >95% and without any N-or C-terminal modifications. The peptides were provided as TFA salts, which were converted to acetate salts, and subsequently dissolved in double-distilled water (ddH_2_O). Short peptides that were difficult to synthesize or purify were excluded from further testing. All producible peptides underwent qualitative solubility assessment and were confirmed to be soluble.

### 3.7. Cell Culture and Senescence Modeling

HEK293 cells were utilized for large-scale peptide screening. Although HEK293 is an immortalized cell line, it remains a valid model for senescence induction due to the following reasons: (1) it retains functional p53 and Rb pathways, which are central to stress-induced senescence; (2) it exhibits well-documented responses to oxidative and genotoxic stressors, leading to stable senescence phenotypes; and (3) its high reproducibility and ease of culture make it suitable for high-throughput screening.

Cells were maintained in Dulbecco’s Modified Eagle Medium (DMEM) supplemented with 10% fetal bovine serum (FBS) and 1% penicillin–streptomycin solution. All cultures were kept in a humidified incubator at 37 °C with 5% CO_2_.

For senescence induction, HEK293 cells were seeded in 96-well plates at a uniform density. Two treatment paradigms were employed 1) Co-treatment: Cells were exposed to 300 μM hydrogen peroxide (H_2_O_2_) for 24 h, followed by recovery in complete medium for another 24 h. Peptides were present throughout both the H_2_O_2_ treatment and recovery phases; 2) Post-treatment: Cells were treated with 300 μM H_2_O_2_ for 24 h, recovered in complete medium for 24 h, and then exposed to peptides for an additional 24 h.

Each peptide was tested at concentrations of 1 μM, 10 μM, and 100 μM. All conditions were performed in triplicate, and the entire experiment was independently repeated three times.

### 3.8. Cell Senescence and Functional Assays

#### 3.8.1. Senescence-Associated β-Galactosidase (SA-β-gal) Staining

SA-β-gal activity was measured using a commercially available kit. Cells were fixed with the provided fixative, washed with PBS, and incubated with β-galactosidase substrate working solution (pH 6.0) at 37 °C under CO_2_-free conditions in the dark until sufficient blue precipitate formed. The reaction was stopped, and the absorbance at 605 nm (A6_05_) was recorded using a microplate reader.

Wells without substrate or cells served as background controls. Since the insoluble indigo precipitate correlates with SA-β-gal activity, lower A6_05_ values indicate reduced senescence.

#### 3.8.2. Intracellular Reactive Oxygen Species (ROS) Measurement

ROS levels were detected using the fluorescent probe DCFH-DA. After incubation, esterase activation, and washing according to the manufacturer’s instructions, the relative fluorescence intensity was measured at excitation/emission wavelengths of 485/535 nm using a multifunctional microplate reader. All steps were performed under consistent incubation times and protected from light to prevent photobleaching.

#### 3.8.3. Mitochondrial Membrane Potential (ΔΨm) Assessment

ΔΨm was evaluated using Mito-Tracker Red CMXRos dye (Beyotime Biotechnology). Freshly prepared dye was applied, followed by incubation and PBS washes. Fluorescence was measured at appropriate excitation/emission wavelengths. To correct for variations in cell number or dye loading, signals were normalized to nuclear dye fluorescence or total protein content from the same or parallel wells. Alternatively, data were normalized to the model control group within each plate during statistical analysis.

### 3.9. *C. elegans* Strains and Maintenance

The N2 strain served as the wild-type control in all experimental procedures. Animals were cultured under standard conditions at 20°C on nematode growth medium (NGM) plates containing 25 mM NaCl, 1.7% agar, 2.5 mg/mL peptone, 5 μg/mL cholesterol, 1 mM CaCl_2_, 1 mM MgSO_4_, and 50 mM KH_2_PO_4_ (pH 6.0).

To eliminate potential metabolic interference from live bacteria, a uniform food source consisting of UV-killed *Escherichia coli* OP50 was utilized throughout the study. Bacterial lawns were prepared by spreading concentrated OP50 suspension onto NGM plates, air-drying briefly, and subjecting to UV irradiation at 999,900 μJ/cm^2^ for 12 minutes using a UV crosslinker with lids removed.

Population synchronization was achieved through hypochlorite treatment. Gravid adults were collected with M9 buffer (22 mM KH_2_PO_4_, 42 mM Na_2_HPO_4_, 86 mM NaCl, 1 mM MgSO_4_) and exposed to alkaline hypochlorite solution (1% NaClO, 0.5 M NaOH) for 10 minutes with continuous agitation. After egg release, samples were centrifuged at 1000 × g for 1 min, washed three times with M9 buffer, and allowed to hatch overnight in M9 at 20°C. Resulting L1 larvae were transferred to OP50-seeded NGM plates for development under controlled temperature conditions.

## 4. Conclusion

In this study, we developed ElixirSeeker2, the first computational framework for *de novo* design of anti-aging peptides that integrates activity prediction, safety constraints, and large-scale virtual screening. Guided by the Information Theory of Aging, ElixirSeeker2 successfully identified multiple short peptides capable of delaying or reversing senescence-associated phenotypes in vitro and enhancing locomotor function in *C. elegans*. These findings demonstrate that systematic *de novo* design can yield functional anti-aging candidates beyond the limits of naturally evolved sequences, providing a generalizable paradigm for next-generation biologics.

It should be emphasized that the present preprint represents only the first phase of a long-term project. We are currently conducting evaluations in rat models to further validate the physiological efficacy and safety of the leading peptides. In parallel, the ElixirSeeker2 algorithm will be further optimized, incorporating advanced structural predictors. Together, these ongoing efforts aim to transform ElixirSeeker2 from a proof-of-concept computational platform into a fully translational framework for anti-aging peptide therapeutics.

## Ethics approval and consent to participate

Not applicable.

## Consent for publication

Not applicable.

## Author Contributions

Conceptualization, B.X. and Y.P.; methodology, Y.P., L.F., N.Z. and H.C.; software, Y.P. and J.Z.(Jingyuan Zhu); validation, Z.J., J.Z.(Jiexin Zheng), L.F., N.Z., H.M. and W.Z.; formal analysis, J.Z.(Jiexin Zheng), W.G., W.X. and R.L.(Ruofei Li); investigation, Y.P., B.X., and S.G.; resources, B.X., N.J., J.H., S.G. and G.L.; data curation, Y.P.; writing—original draft preparation, Y.P. and H.C.; writing—review and editing, F.Y., Y.P., J.Z.(Jingyuan Zhu), G.L. and B.X.; visualization, R.L.(Rui Liang), R.L.(Ruofei Li) and W.X.; supervision, S.G. and B.X.; project administration, B.X., J.N. and S.G.; funding acquisition, S.G., B.X., J.N. and G.L. All authors have read and agreed to the published version of the manuscript.

## Acknowledgments

Special thanks to Dr. Yuxuan Lyv for their invaluable advice and Ms. Nature Belle for her assistance. We also extend our appreciation to Aging laboratory technicians for their diligent work in maintaining the *C. elegans* cultures and to the administrative staff for their support throughout the project. Furthermore, we would like to express our gratitude to UESTC_BioMed for their revisions of the figures and language in this manuscript.

---

**Table S1.**
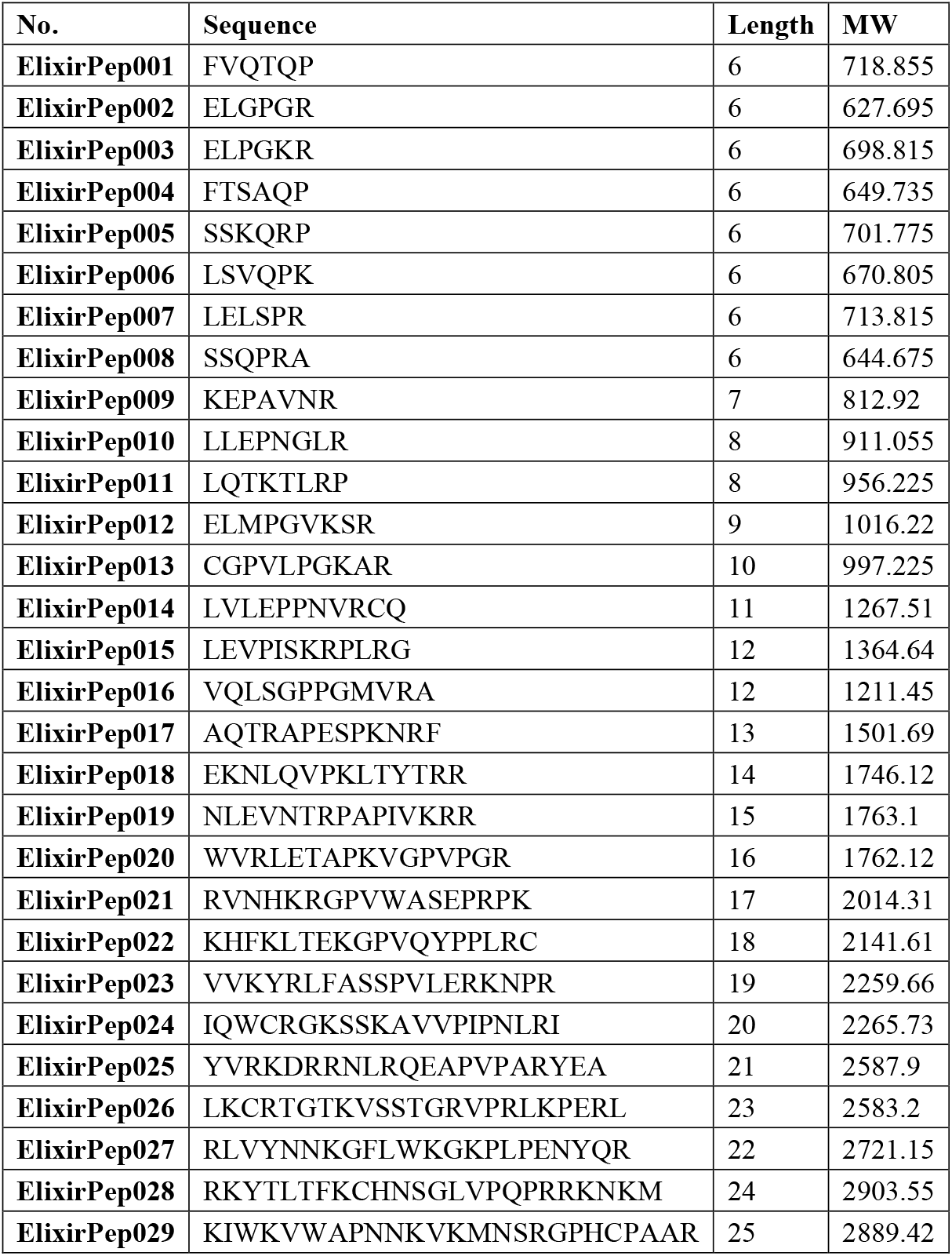
Candidates’ Sequence.

**Figure S1.**
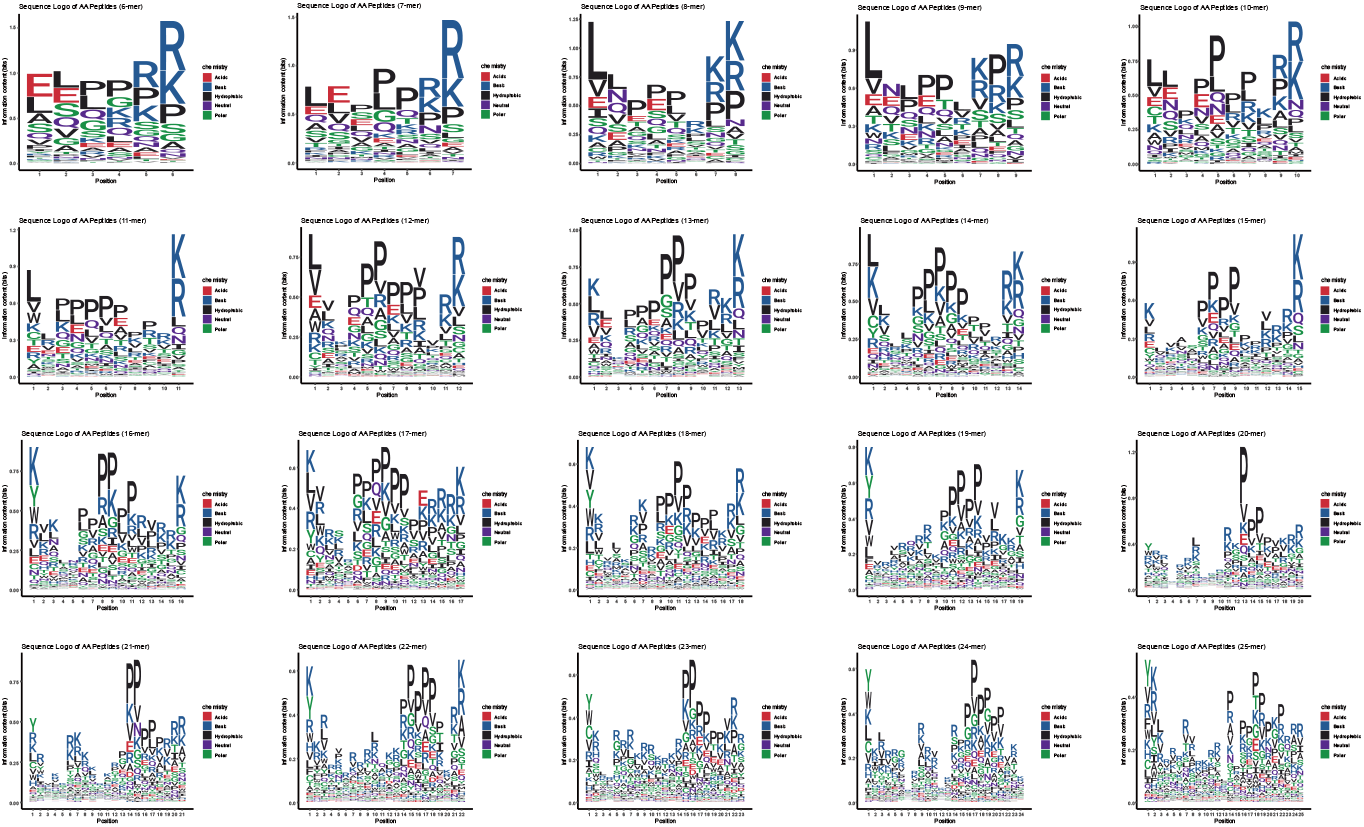
Seqlogo of each length of AAPeptide.

## Reference

[1] López-Otín C, Blasco MA, Partridge L, Serrano M, Kroemer G. Hallmarks of aging: An expanding universe. Cell. 2023 Jan 19;186(2):243–278. doi: 10.1016/j.cell.2022.11.001. Epub 2023 Jan 3. PMID: 36599349.

[2] Melo Dos Santos LS, Trombetta-Lima M, Eggen B, Demaria M. Cellular senescence in brain aging and neurodegeneration. Ageing Res Rev. 2024 Jan;93:102141. doi: 10.1016/j.arr.2023.102141. Epub 2023 Nov 27. PMID: 38030088.

[3] Moskalev A, Guvatova Z, Lopes IA, Beckett CW, Kennedy BK, De Magalhaes JP, Makarov AA. Targeting aging mechanisms: pharmacological perspectives. Trends Endocrinol Metab. 2022 Apr;33(4):266–280. doi: 10.1016/j.tem.2022.01.007. Epub 2022 Feb 17. PMID: 35183431.

[4] Semsei I. On the nature of aging. Mech Ageing Dev. 2000 Aug 15;117(1-3):93–108. doi: 10.1016/s0047-6374(00)00147-0. PMID: 10958926.

[5] Duan R, Fu Q, Sun Y, Li Q. Epigenetic clock: A promising biomarker and practical tool in aging. Ageing Res Rev. 2022 Nov;81:101743. doi: 10.1016/j.arr.2022.101743. Epub 2022 Oct 4. PMID: 36206857.

[6] la Torre A, Lo Vecchio F, Greco A. Epigenetic Mechanisms of Aging and Aging-Associated Diseases. Cells. 2023 Apr 14;12(8):1163. doi: 10.3390/cells12081163. PMID: 37190071; PMCID: PMC10136616.

[7] Li A, Koch Z, Ideker T. Epigenetic aging: Biological age prediction and informing a mechanistic theory of aging. J Intern Med. 2022 Nov;292(5):733–744. doi: 10.1111/joim.13533. Epub 2022 Jun 20. PMID: 35726002.

[8] Lu YR, Tian X, Sinclair DA. The Information Theory of Aging. Nat Aging. 2023 Dec;3(12):1486–1499. doi: 10.1038/s43587-023-00527-6. Epub 2023 Dec 15. PMID: 38102202.

[9] Yang JH, Petty CA, Dixon-McDougall T, Lopez MV, Tyshkovskiy A, Maybury-Lewis S, Tian X, Ibrahim N, Chen Z, Griffin PT, Arnold M, Li J, Martinez OA, Behn A, Rogers-Hammond R, Angeli S, Gladyshev VN, Sinclair DA. Chemically induced reprogramming to reverse cellular aging. Aging (Albany NY). 2023 Jul 12;15(13):5966–5989. doi: 10.18632/aging.204896. Epub 2023 Jul 12. PMID: 37437248; PMCID: PMC10373966.

[10] Pereira B, Correia FP, Alves IA, Costa M, Gameiro M, Martins AP, Saraiva JA. Epigenetic reprogramming as a key to reverse ageing and increase longevity. Ageing Res Rev. 2024 Mar;95:102204. doi: 10.1016/j.arr.2024.102204. Epub 2024 Jan 23. PMID: 38272265.

[11] Lu Y, Brommer B, Tian X, Krishnan A, Meer M, Wang C, Vera DL, Zeng Q, Yu D, Bonkowski MS, Yang JH, Zhou S, Hoffmann EM, Karg MM, Schultz MB, Kane AE, Davidsohn N, Korobkina E, Chwalek K, Rajman LA, Church GM, Hochedlinger K, Gladyshev VN, Horvath S, Levine ME, Gregory-Ksander MS, Ksander BR, He Z, Sinclair DA. Reprogramming to recover youthful epigenetic information and restore vision. Nature. 2020 Dec;588(7836):124–129. doi: 10.1038/s41586-020-2975-4. Epub 2020 Dec 2. PMID: 33268865; PMCID: PMC7752134.

[12] Buchanan S, Combet E, Stenvinkel P, Shiels PG. Klotho, Aging, and the Failing Kidney. Front Endocrinol (Lausanne). 2020 Aug 27;11:560. doi: 10.3389/fendo.2020.00560. PMID: 32982966; PMCID: PMC7481361.

[13] Zeng C, Chen M. Progress in Nonalcoholic Fatty Liver Disease: SIRT Family Regulates Mitochondrial Biogenesis. Biomolecules. 2022 Aug 5;12(8):1079. doi: 10.3390/biom12081079. PMID: 36008973; PMCID: PMC9405760.

[14] Martins R, Lithgow GJ, Link W. Long live FOXO: unraveling the role of FOXO proteins in aging and longevity. Aging Cell. 2016 Apr;15(2):196–207. doi: 10.1111/acel.12427. Epub 2015 Dec 8. PMID: 26643314; PMCID: PMC4783344.

[15] Lim JJ, Noh S, Kang W, Hyun B, Lee BH, Hyun S. Pharmacological inhibition of USP14 delays proteostasis-associated aging in a proteasome-dependent but foxo-independent manner. Autophagy. 2024 Dec;20(12):2752–2768. doi: 10.1080/15548627.2024.2389607. Epub 2024 Aug 15. PMID: 39113571; PMCID: PMC11587835.

[16] Ma B, Liu D, Wang Z, Zhang D, Jian Y, Zhang K, Zhou T, Gao Y, Fan Y, Ma J, Gao Y, Chen Y, Chen S, Liu J, Li X, Li L. A Top-Down Design Approach for Generating a Peptide PROTAC Drug Targeting Androgen Receptor for Androgenetic Alopecia Therapy. J Med Chem. 2024 Jun 27;67(12):10336–10349. doi: 10.1021/acs.jmedchem.4c00828. Epub 2024 Jun 5. PMID: 38836467.

[17] Erak M, Bellmann-Sickert K, Els-Heindl S, Beck-Sickinger AG. Peptide chemistry toolbox - Transforming natural peptides into peptide therapeutics. Bioorg Med Chem. 2018 Jun 1;26(10):2759–2765. doi: 10.1016/j.bmc.2018.01.012. Epub 2018 Jan 31. PMID: 29395804.

[18] Paulus J, Sewald N. Small molecule- and peptide-drug conjugates addressing integrins: A story of targeted cancer treatment. J Pept Sci. 2024 Jul;30(7):e3561. doi: 10.1002/psc.3561. Epub 2024 Feb 21. PMID: 38382900.

[19] Yuan Q, Ren Q, Li L, Tan H, Lu M, Tian Y, Huang L, Zhao B, Fu H, Hou FF, Zhou L, Liu Y. A Klotho-derived peptide protects against kidney fibrosis by targeting TGF-β signaling. Nat Commun. 2022 Jan 21;13(1):438. doi: 10.1038/s41467-022-28096-z. Erratum in: Nat Commun. 2022 Nov 4;13(1):6640. doi: 10.1038/s41467-022-34454-8. PMID: 35064106; PMCID: PMC8782923.

[20] Kortemme T. De novo protein design-From new structures to programmable functions. Cell. 2024 Feb 1;187(3):526–544. doi: 10.1016/j.cell.2023.12.028. PMID: 38306980; PMCID: PMC10990048.

[21] Pan Y, Cai H, Ye F, Xu W, Huang Z, Zhu J, Gong Y, Li Y, Ezemaduka AN, Gao S, Liu S, Li G, Li H, Yang J, Ning J, Xian B. ElixirSeeker: A Machine Learning Framework Utilizing Fusion Molecular Fingerprints for the Discovery of Lifespan-Extending Compounds. Aging Cell. 2025 Aug;24(8):e70116. doi: 10.1111/acel.70116. Epub 2025 May 26. PMID: 40419453; PMCID: PMC12341795.

[22] Arora S, Mittal A, Duari S, Chauhan S, Dixit NK, Mohanty SK, Sharma A, Solanki S, Sharma AK, Gautam V, Gahlot PS, Satija S, Nanshi J, Kapoor N, Cb L, Sengupta D, Mehrotra P, Ghosh TS, Ahuja G. Discovering geroprotectors through the explainable artificial intelligence-based platform AgeXtend. Nat Aging. 2025 Jan;5(1):144–161. doi: 10.1038/s43587-024-00763-4. Epub 2024 Dec 3. PMID: 39627462.

[23] Gainza P, Wehrle S, Van Hall-Beauvais A, Marchand A, Scheck A, Harteveld Z, Buckley S, Ni D, Tan S, Sverrisson F, Goverde C, Turelli P, Raclot C, Teslenko A, Pacesa M, Rosset S, Georgeon S, Marsden J, Petruzzella A, Liu K, Xu Z, Chai Y, Han P, Gao GF, Oricchio E, Fierz B, Trono D, Stahlberg H, Bronstein M, Correia BE. De novo design of protein interactions with learned surface fingerprints. Nature. 2023 May;617(7959):176–184. doi: 10.1038/s41586-023-05993-x. Epub 2023 Apr 26. PMID: 37100904; PMCID: PMC10131520.

[24] Bennett NR, Coventry B, Goreshnik I, Huang B, Allen A, Vafeados D, Peng YP, Dauparas J, Baek M, Stewart L, DiMaio F, De Munck S, Savvides SN, Baker D. Improving de novo protein binder design with deep learning. Nat Commun. 2023 May 6;14(1):2625. doi: 10.1038/s41467-023-38328-5. PMID: 37149653; PMCID: PMC10163288.

[25] Pillai A, Idris A, Philomin A, Weidle C, Skotheim R, Leung PJY, Broerman A, Demakis C, Borst AJ, Praetorius F, Baker D. De novo design of allosterically switchable protein assemblies. Nature. 2024 Aug;632(8026):911–920. doi: 10.1038/s41586-024-07813-2. Epub 2024 Aug 14. PMID: 39143214; PMCID: PMC11338832.

[26] Mallik BB, Stanislaw J, Alawathurage TM, Khmelinskaia A. De Novo Design of Polyhedral Protein Assemblies: Before and After the AI Revolution. Chembiochem. 2023 Aug 1;24(15):e202300117. doi: 10.1002/cbic.202300117. Epub 2023 Jul 12. PMID: 37014094.

[27] Hu Y, Zhou K Y, Wang Z, et al. N-stearoyl-l-Tyrosine inhibits the cell senescence and apoptosis induced by H2O2 in HEK293/Tau cells via the CB2 receptor[J]. Chemico-Biological Interactions, 2017, 272: 135–144.

[28] Zhang H, Cui N, Ma X, et al. Structural basis of augmenting taurine uptake by the taurine transporter in alleviating cellular senescence[J]. Cell Research, 2025: 1–4.

[29] Zhou J, Liu K, Bauer C, et al. Modulation of cellular senescence in HEK293 and HepG2 cells by Ultrafiltrates UPla and ULu is partly mediated by modulation of mitochondrial homeostasis under oxidative stress[J]. International Journal of Molecular Sciences, 2023, 24(7): 6748.

